# Responses of Oil Palm Pollinator, *Elaeidobius kamerunicus* to Different Concentrations of Estragoles

**DOI:** 10.1101/2021.03.24.436726

**Authors:** Muhammad Fahmi-Halil, Mohamad Haris-Hussain, Razali Mirad, AB Idris, Johari Jalinas

## Abstract

*Elaeidobius kamerunicus* is the main insect pollinator for oil palm (*Elaeis guineensis*) worldwide. One of the main reason *E. kamerunicus* attracted to oil palm inflorescences is estragole, a volatile organic compound released by the oil palm inflorescences during anthesis stage. However, the amount of estragole released from the oil palm inflorescence is varied due to the influence of abiotic and biotic factors and is seen to have an impact on *E. kamerunicus* pollination activity on the oil palm. To evaluate the responses of *E. kamerunicus*, different types (wild and reared) and sex (male and female) of *E. kamerunicus* were exposed to different concentrations (1, 5, 10, 30, 50, 70, 100, 150 and 200 ppm) of commercial estragole using four-arm olfactometer. Results showed that *E. kamerunicus* significantly preferred 100 ppm of estragole compared to other concentration (F = 139.81; d.f. = 9; P < 0.05). A significant interaction was also recorded between estragole concentrations and sexes of *E. kamerunicus* (F = 3.91; d.f. = 9; P < 0.05) where male *E. kamerunicus* was found to be more responsive to 100 ppm of estragole compared female *E. kamerunicus*. The *E. kamerunicus* responses to estragole is in line with the increase of estragole concentration up to 100 ppm. However, the response of *E. kamerunicus* was significantly decreased after the concentration value. The result of this study can be a good platform for future references since the estragole compound plays a significant role in oil palm’s flower pollination by *E. kamerunicus*. The factor of type and sexes of *E. kamerunicus* did not affect the preferences which indicated that *E. kamerunicus* reared in the laboratory have the potential to be released into the oil palm plantation area to overcome the problem of pollination.

## INTRODUCTION

Oil palm is one of the important commodity crops that highly dependent on insect pollinators for reproduction. Interestingly, this plant has a special insect pollinator (*Elaeidobius kamerunicus*) that only pollinates the oil palm inflorescence (Setyamidjaja, 2006; Sambathkumar & Ranjith, 2011; Zahari et al., 2019). It was brought from Cameroon to Malaysia and other countries in the early 80s for release into the oil palm plantations (Caudwell et al., 2003; Appiah & Agyei, 2013). The presence of *E. kamerunicus* pollinator has successfully solved the pollination problem encountered in oil palm plantations over time and has led to an increase in oil production in most countries such as Malaysia, Indonesia and India (Ponnamma, 1999; Caudwell et al., 2003).

One of the factors that causes *E. kamerunicus* pollinators to be attracted to oil palm is the suitability of palm flowers to serve as a habitat and food source for the insects. The life cycle of the *E. kamerunicus* pollinator occurs in the oil palm flower where the eggs, larvae and pupae live inside of the male flower. The adult weevil lives by feeding and mating around the outside of the flower (Tuo et al., 2011). The presence of *E. kamerunicus* is also driven by the odor factor of palm flowers released during the flower anthesis stage. Recent research shows that *E. kamerunicus* pollen tend to prefer oil palm over other species of palm flowers because they are attracted to the aromas and odors produced by the oil palm (Corley & Tinker, 2003; Adaigbe et al., 2014). Whilst, chemical studies on oil palm flowers have found that the odor produced by these flowers is due to the volatile compounds of estragole (Anggraeni et al., 2013; Fahmi et al., 2016).

Estragole is one of the volatile organic compounds (VOC) most found in plants, especially in herbal and aromatic plants (Raffo et al., 2011; Yamani et al., 2014). It acted as insect attractants for several types of flowering plants such as *Cycas revoluta*, *Hyssopus officinalis, Agastache rugosa* including palm oil (Hiroshi & Masumi, 2006; Leslie & Richard 2004; Tandon et al., 2001).

Although estragole compound have been identified as one of the key factors of *E. kamerunicus* pollinator attraction to oil palm trees (Hussein et al., 1991; Appiah & Agyei-Dwarko, 2013), studies of the effects of this compound on weevil behaviour are still less and poorly conducted. Most of *E. kamerunicus* pollinator studies are more focused on its life cycle and the distribution of this weevil population in the oil palm plantations. Thus, a study was conducted in the laboratory using commercial estragole compound to understand the role of this compound in its interaction with *E. kamerunicus* pollinator. The objective of this study was to determine response of the type (wild and reared) and sex of *E. kamerunicus* pollinator to different concentrations of estragole compounds.

## MATERIALS AND METHODS

### Preparation of *Elaeidobius kamerunicus* adult sample

Study was conducted at the biological control laboratory, Agrobiodiversity and Environment Research Center, MARDI Serdang. The *E. kamerunicus* samples used in this study were of Laboratory-Reared and Field-Collected. Sample *E. kamerunicus* was obtained from MPOB Keratong oil palm plantation, Pahang, Malaysia. Sampling was done by packing mature male flowers (end stage anthesis) on palm trees using gauze cloth (Zahari et al. 2019). A total of 1000 samples of these field-collected *E. kamerunicus* were brought to the laboratory and kept at room temperature around 26-28°C and 12:12 (light:dark) where half of the samples were used in this experiment and others were kept for breeding to obtain laboratory-reared type of *E. kamerunicus* (first generation).

A total of 500 individuals of male and female *E. kamerunicus* were placed in several plastic container (16 cm Height × 10 cm Width diameter) for rearing. Each of the plastic containers was provided with five male flower spikelets (anthesis) as food sources and breeding sites. After four days, the spikelets were removed and transferred into another container and left at room temperature until the emergence of new *E. kamerunicus* laboratory-reared adults from the spikelets.

### Preparation of estragole compound solution

The standard estragole compounds used in this study were bought from the company Sigma-Aldrich, USA. A total of nine series of estragole concentrations were prepared in this study at 1, 5, 10, 30, 50, 70, 100, 150 and 200 μg / mL where each series was mixed with hexane solvent (RCI Labscan) to obtain a 10 mL mixed solution. To obtain these series of different estragole concentrations, the standard dilution method (M_1_V_1_= M_2_V_2_) was used.

### Olfactometric bioassays

Experiment was conducted by using a four-arm olfactometer with a square shape of 107 cm (length) × 107 cm (width) (Toption, China) (Fig. 1). The olfactometer was divided into two parts. The main part was made of white hard acrylic fiber and covered with clear thick plastic on top. It consisted of four arms with four opening holes and one hole in the middle. Each hole opening was fitted with a circular glass flanged with the end of which has a small open channel for airflow. A glass cylinder containing charcoal powder was used to absorb the environment lab odors. The hole in the middle of the device was connected to the suction pump for circulation of air flow inside the olfactometer section during the experiment.

**Fig. 1.**
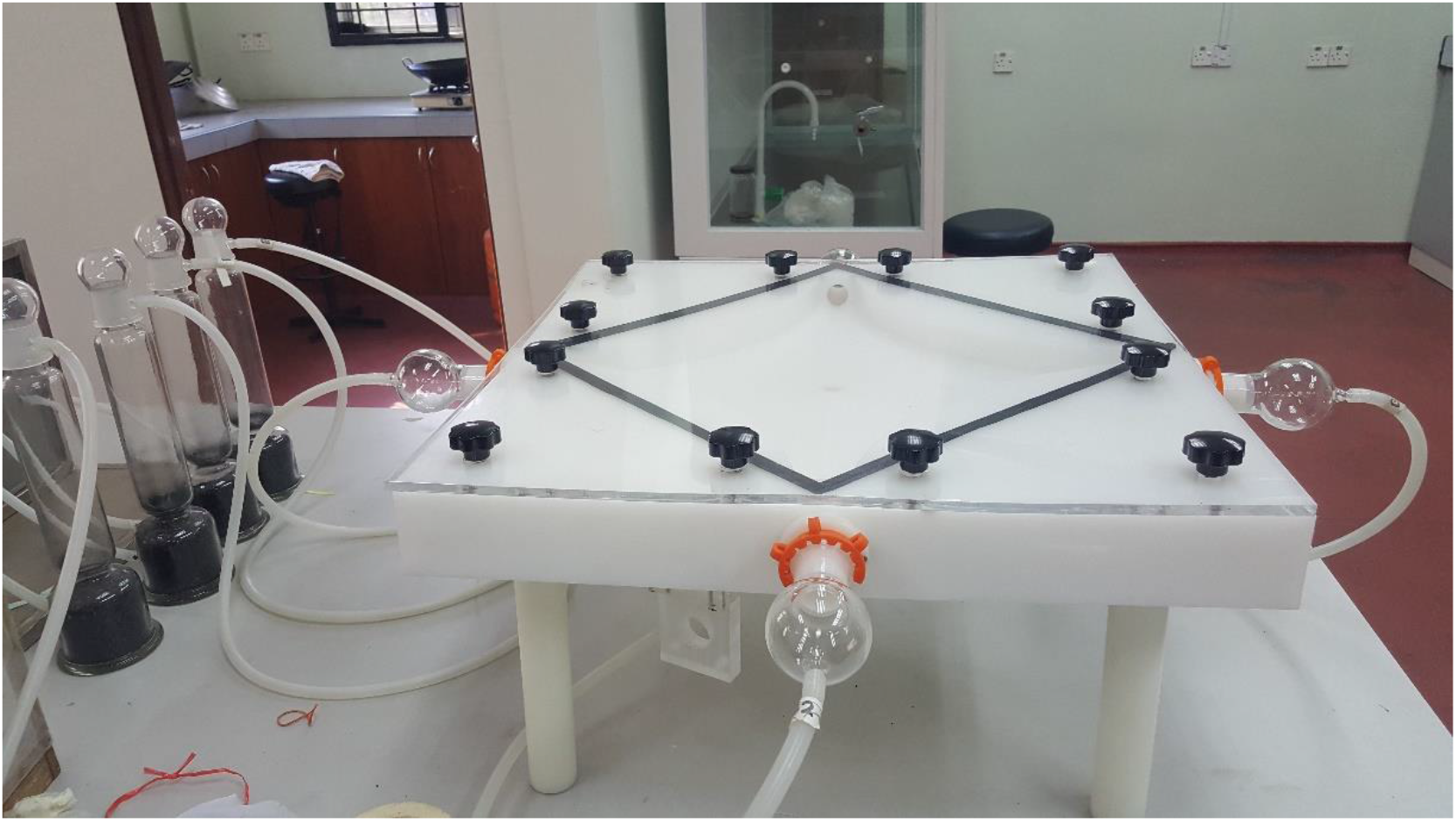
Four arm olfactometer with glass flask at the end of each arm.

Experimental methods were based on the olfactometer study conducted by Haris-Hussain et al. (2020) with some modifications. A total of 900 *E. kamerunicus* adults comprising of field-collected (450 individuals) and laboratory-reared (450 individuals) types were used in this study. This experiment was conducted from 09.00 to 17.00 hrs in the biological control laboratory with room temperature of 25-27°C, 60-80% relative humidity (RH) and in illuminated conditions using fluorescent lamps throughout the experiment. Behavioral assessment for both *E. kamerunicus* type was conducted separately. Each weevil tested was isolated 12 hours prior the start of the bioassay experiments. Each assay began by releasing a total of 10 weevils consisting of five males and females at the center of the olfactory equipment. Each side of this olfactometer was fitted with a round glass flask filled with cotton that diluted in different estragole concentration solution (1 ml). *E. kamerunicus* adults in each assay were exposed to three different series of estragole concentrations (concentrations of 1 ppm, 5 ppm, 10 ppm, control; 30 ppm, 50 ppm, 70 ppm, control; 100 ppm, 150 ppm, 200 ppm, control) with control treatment (no estragole) in each series concentration. Experiments for each series of estragole concentrations were repeated 15 times (replicates).

Number of individual *E. kamerunicus* responded to estragoles given in olfactormter was counted. The time given for each assay was 30 minutes. Different *E. kamerunicus* was used once for each experiment. After every assay was completed, the olfactometer’s cover was opened and the glass flasks were swapped with alcohol to extinguish the trapped odour before starting new assay.

### Data analyses

To get the percentage of *E. kamerunicus* responded to each treatment, the number of individual *E. kamerunicus* responded divided by the total number of *E. kamerunicus* multiply with 100%. Percentage of *E. kamerunicus* responses towards different estragole concentrations were then transformed using arcsine transformation (ASIN√x) for statistical analysis. Three-way Analysis of Variance (ANOVA) and Tukey’s mean separation tests were used to analyzed the response of *E. kamerunicus* (sexes, types) to different estragole concentrations. Turkey’s test was used to separate the treatment means at P < 0,05 in Minitab 16.0.

## RESULTS

### Interactions among sex, type and estragole concentration factors on *E. kamerunicus* preferences

A three-way ANOVA analysis showed there was no significant interaction among factors (*E. kamerunicus* sex, type and estragole concentrations) (ANOVA, p > 0.05) (Table 1). Only concentration (F = 139.81; d.f. = 9; p < 0.05) and sex*concentrations (F = 3.91; d.f. = 9; p < 0.05) showed a significant different.

**Table 1.** THREE-WAY ANOVA RESULTS FOR THE STUDY OF *E. kamerunicus* POLLINATOR SUSCEPTIBILITY TO DIFFERENT CONCENTRATIONS OF ESTRAGOLE COMPOUNDS.

| Source | Df | F-value | P-value |
| --- | --- | --- | --- |
| Type of *E. kamerunicus* | 1 | 2.27 | > 0.05 |
| Sex of *E. kamerunicus* | 1 | 0.07 | > 0.05 |
| Concentration of estragole | 9 | 139.81 | < **0.05** |
| Type * Sex | 1 | 0.24 | > 0.05 |
| Type * Concentration | 9 | 0.96 | > 0.05 |
| Sex * Concentration | 9 | 3.91 | < **0.05** |
| Type * Sex * Concentration | 9 | 0.93 | > 0.05 |
| Error | 1160 |  |  |
| Total | 1199 |  |  |

### Responses of *E. kamerunicus* to different concentrations of estragole

Result shows that *E. kamerunicus* responded significantly more (49.5% ± 1.7) significantly responded toward a concentration of 100 ppm of estragole compared to other concentration (p < 0.05), (Fig. 2). Interestingly, the mean percentage of *E. kamerunicus* adult individual responded significantly less to 150 and 200 ppm estragole concentrations. Similar result was observed between 150 ppm and lower estragole concentrations (p > 0.05), similar with between 30 ppm, 10 ppm and 5 ppm of estragole concentration. The lowest mean percentage of *E. kamerunicus* was recorded at 1 ppm (3.1% ± 0.7) and significantly different (p<0.05) among other estragole concentration except with 5 ppm. However, there were no *E. kamerunicus* individual recorded in control treatment during this study.

**Fig. 2.**
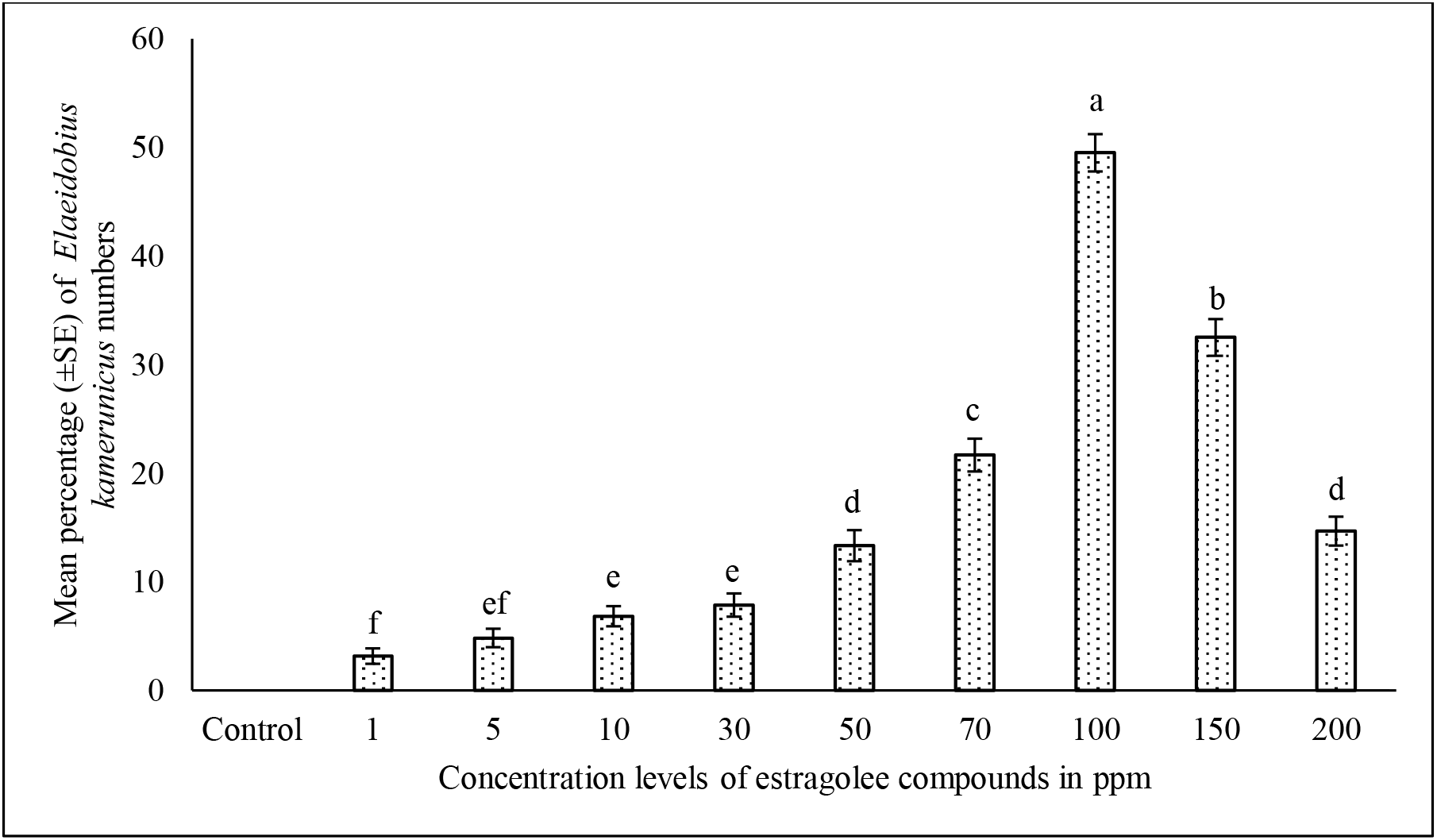
Min percentage (± SE) number of weevil *E. kamerunicus* recorded between different concentrations of estragole compounds.

### Interaction between different sex and concentration factors on *E. kamerunicus* preferences

The interaction between *E. kamerunicus* sexes and estragole concentrations showed that male and female of *E. kamerunicus* recorded a significant high mean percentage of weevils preferences at 100 ppm estragole concentration (55.6% ± 2.5) and (43.3% ± 2.0) respectively compared to other concentrations (p < 0.05) (Fig. 3). The lowest mean percentage of weevils responded by the weevil was at 1 ppm estragole concentration (3.3% ± 1.0) and (3.0%. ± 0.9 for male and female,) respectively.

**Fig. 3.**
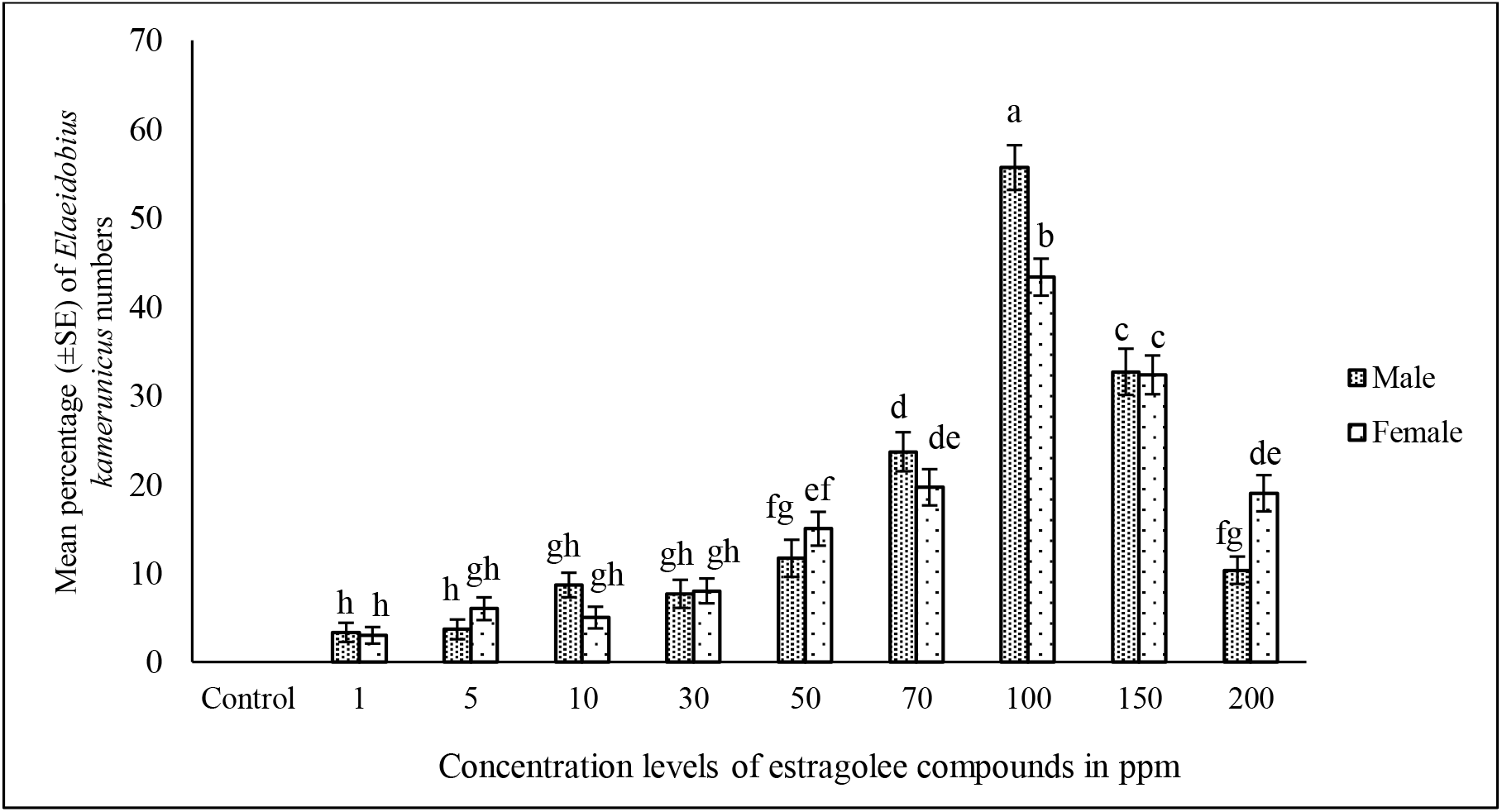
Min percentage (± SE) of the number of weevil *E. kamerunicus* recorded as a result of interaction between weevil sexes with different concentration of estragole compounds.

The interaction results also showed a significant difference between mean percentage of male and female of *E. kamerunicus* at 200 ppm estragole concentration. However, there was no significant difference (p > 0.05) of mean percentage between male and female of *E. kamerunicus* at 150 ppm, 70 ppm, 50 ppm, 30 ppm, 5 ppm and 1 ppm estragole concentrations.

### Effect of types and sexes on *E. kamerunicus* responses to various concentrations of estragole

There was no significant difference in the number of *E. kamerunicus* individual responded to estragole concentrations (Fig. 4). Similar result was also observed for types of *E. kamerunicus* (Fig. 5). While for the sex factors of *E. kamerunicus* (Figure 5).

**Fig. 4.**
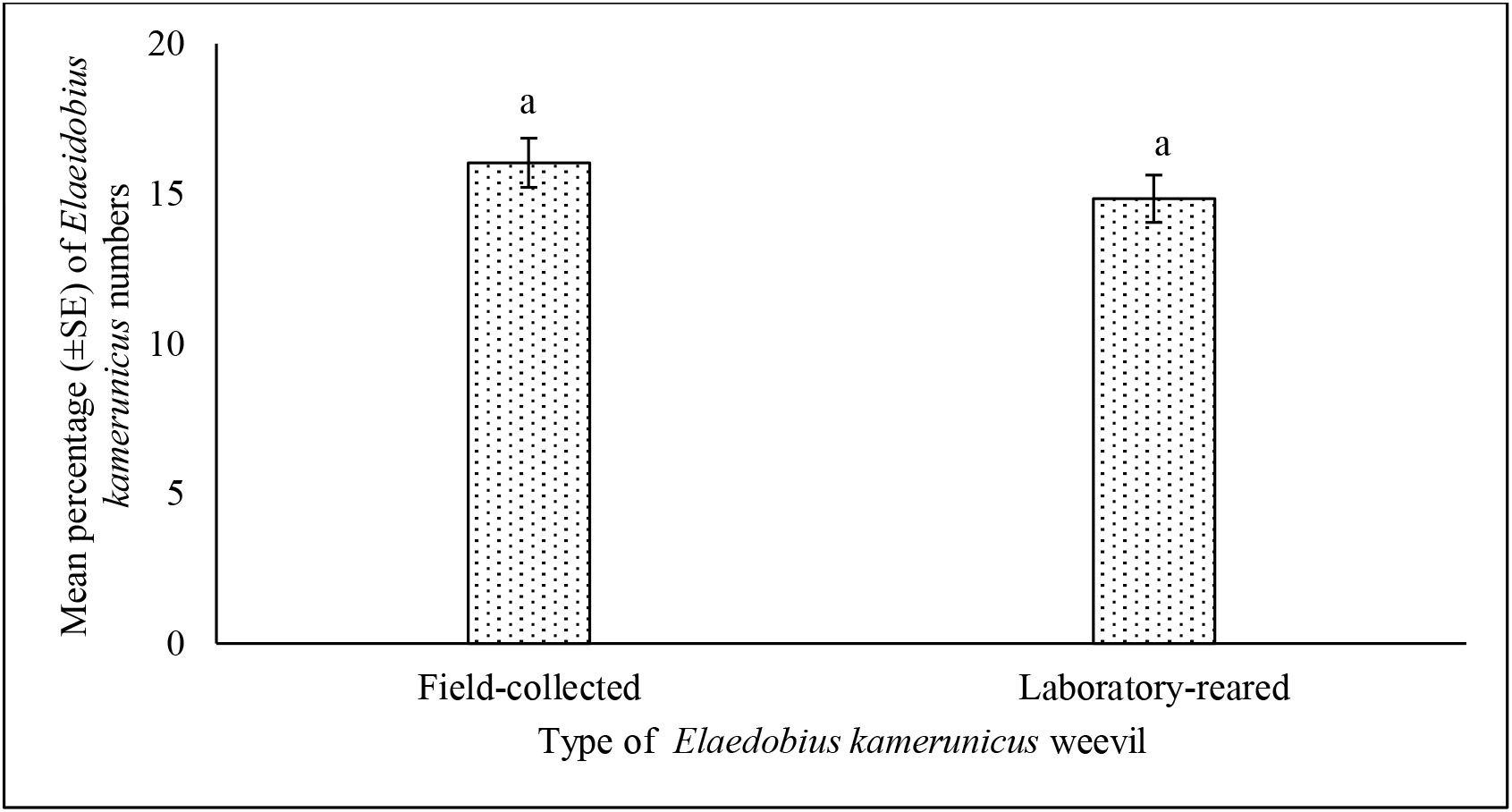
Min percentage (± SE) number of weevils between field-collected and laboratory-reared type of *E. kamerunicus*

**Fig. 5.**
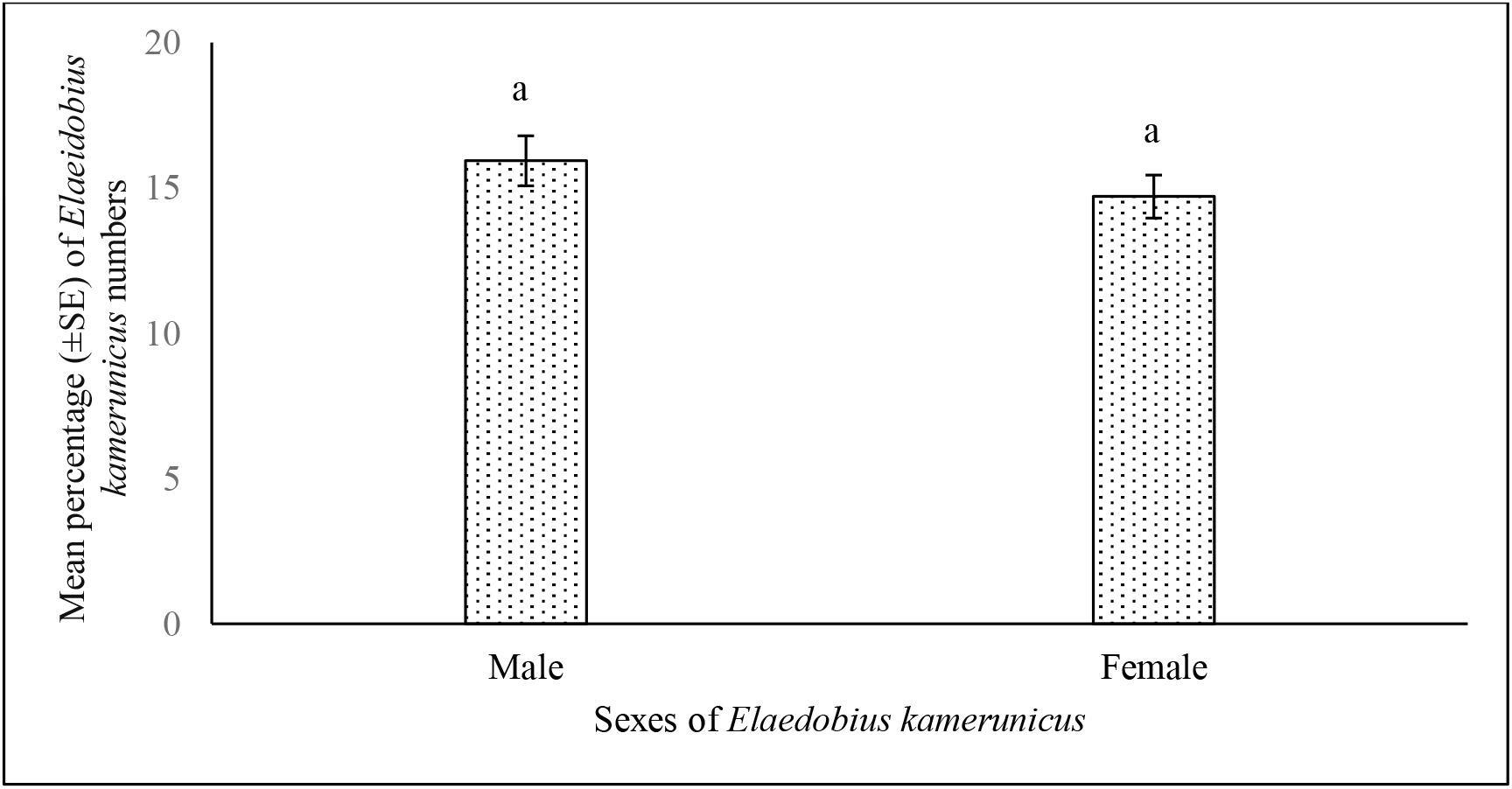
Min percentage (± SE) number of weevils between *E. kamerunicus* male and female

## DISCUSSION

Overall, this study found that *E. kamerunicus* was attracted to estragole compounds but varied with estragole concentrations tested. These findings are in par with previous studies showing that estragole compound is one of the key factors for *E. kamerunicus* to response and attracted to oil palm inflorescence most probably due to the compound has a strong aroma that *E. kamerunicus* prefers (Hazimah, 1990; Hussein et al., 1991; Anggraeni et al., 2013; Tandon et al., 2001). This aroma is derived from a benzene ring found in the structure of estragole, which is found mainly in aromatic plants such as *Artemisia dracunculus* L., *Ocimum basilicum* L., and *Agastache rugosa* (Vincenzi et al. 2000). The results also showed that estragole compound plays an important role in palm flowering activities and its ability of the compound to attract the presence of *E. kamerunicus* although no food source was used. This is the main reason why female flower is able to attract *E. kamerunicus* by releasing estragole compound during the antesis stage thus allow pollination to take place (Sambathkumar & Ranjith, 2011).

The releasing rate of plant-derived organic compounds is strongly influenced by abiotic and biotic factors (Holopainen & Gershenzon, 2010). This condition is thought to have an effect on the insects’ behaviour, especially those insects that use a sense of smell to detect food and their partner (Haris et al., 2014). Thus, the exposure of *E. kamerunicus* to different concentrations of estragole compound in this study have revealed the optimum concentration of estragole compounds in attracting the weevil. The higher and lower than the 100 ppm concentration of estragole compund only lead to low *E. kamerunicus* attractions toward the oil palm flowers, probably because the weevil’s sensory function is impaired by high concentration of the compound. According to Wood (1982), this condition is called ‘multiple function’ in which the sensory function of the insect is under stress and disturbance. This is based on a study of verbenone exposed to bark weevil, *Dendroctonus ponderosae* which, in a high concentration of compound, makes it difficult and confusing to the weevil’s senses to detect this plant-derived volatile compound.

In addition, high concentrations of volatile organic compounds often occur in plants under stress due to abiotic and biotic factors (Dicke et al., 2009; Holopainen & Gershenzon, 2010). This condition not only affect the physiology of plants but also can influence the behaviour of insects where studies have shown that some of the coleopteran weevil avoid from approaching the plants that release higher concentration of volatile organic compounds (Jermy et al., 1998; Ikonen, 2002; Heil, 2004). This might be related with the ability of insect sense to identify the quality of plants based on VOC released, where unhealthy plant normally produced more VOC than the healthy one (Chittka & Raine, 2006; Schiestl, 2015). However, further studies need to be carried out in the field to observe the effect of plant-derived organic compounds on insect’s behaviour.

Besides, type of *E. kamerunicus* was proved to be a non-significant factor in affecting the *E. kamerunicus* response towards estragole concentrations in this study. This probably due to the laboratory-reared *E. kamerunicus* was the first-generation (F1) of field population where they were not significantly different from that of the field-collected *E. kamerunicus* in genetic or physiological characteristics of the weevil. Richgels & Rollmann (2012) reported that, insects that have the same genetic and physiological characters have the same ability to detect VOC released by plants.

Interestingly, our results also found that the weevil’s sex factor did not significantly influence *E. kamerunicu*s’ response to different concentrations of estragole. This indicates that both sexes of *E. kamerunicus* are able to detect and respond well to estragole compounds as evidenced as they could easily observed on both oil palm inflorescenses. This feature probably makes *E. kamerunicus* the best pollinator for oil palm crops since estragole compounds are the major VOCs produced and released by the palm inflorescenece(Misztal et al. 2014; Muhamad Fahmi et al. 2016). This is tend to agree with Farre-Armengol et al. (2015) where they reported the ability of pollinator insects to detect VOCs produced by flowering plants will help the insects to determine the flower’s location more precisely and rapidly increasing the process of pollination at once.

## CONCLUSION

The importance of estragole in guiding the *E. kamerunicus* to pollinate oil inflorescence is well known since 1980s. However, report on it becoming ineffective pollinators has been documented by many. We believed it could due to many factors and estragoles – *E. kamerunicus* relation could one of them. Result of our study showed that *E. kamerunicus* specifically responded well to 100 ppm estragole concentrations in the laboratory experiment irrespective of sexes of the weevils. As we suggested that in order to make *E. kamerunicus* effective pollinator we must create an environment – biotic and abiotic – that ensure oil palm tree and inflorescence physiologically able to emitted estragoles around that concentrations in the field though several factors may involve like soil condition, RH and temperature. Further studies should be conducted in measuring estragoles emission and *E. kamerunicus* population density per in spikelets in relation to soil, planting materials, RH and temperature.

## Acknowledgements

The authors would like to thank all the subjects for participating in the study.

